# A library of promoter-*gfp* fusion reporters for studying systemic expression pattern of cyclic-di-GMP metabolism-related genes in *Pseudomonas aeruginosa*

**DOI:** 10.1101/2022.06.15.496363

**Authors:** Dejian Liu, Di Wang, Qing Wei, Yu Zhang, Luyan Z Ma

## Abstract

The opportunistic pathogen *Pseudomonas aeruginosa* is an environmental microorganism, which is notorious for its resistance or tolerance to antibiotics due to the formation of biofilms. Cyclic diguanosine monophosphate (c-di-GMP) is a bacterial second messenger that plays critical roles in biofilm formation. *P. aeruginosa* contains 41 genes that encode enzymes to participate in the metabolism of c-di-GMP (biosynthesis or degradation), yet it lacks tools to investigate the systemic expression pattern of those genes. Here, we constructed a promoter-*gfp* transcriptional fusion reporters’ library that consists of 41 reporter plasmids. Each plasmid contains a promoter of corresponding <u>c</u>-di-GMP <u>m</u>etabolism-related (CMR) genes from *P. aeruginosa* PAO1 strain, thus each promoter-Gfp fusion reporter can be used to detect the promotor’ activity as well as the transcription of corresponding gene. The promoters’ activity was tested in *P. aeruginosa* and *Escherichia coli* respectively. Among the 41 genes, the promoter of 26 genes showed activity in both *P. aeruginosa* and *E. coli*. The library was applied to determine the influence of different temperatures, growth media, and sub-inhibitory concentrations of antibiotics on transcriptional profile of the 41 CMR genes in *P. aeruginosa*. The results showed different growth conditions did impact different genes’ transcription, while the promoter’ activity of a few genes kept at the same level under several different growth conditions. In summary, we provided a promoter-*gfp* fusion reporters’ library for systemic monitoring or study of the regulation of CMR genes in *P. aeruginosa* and the functional promoters can also be used as a bio-brick for synthetic biology studies.

**Importance:** The opportunistic pathogen *P. aeruginosa* can cause acute and chronic infections in humans and it is one of main pathogens in nosocomial infections. Biofilm formation is one of most important causes for *P. aeruginosa* to persist in hosts and evade immune and antibiotic attacks. c-di-GMP is an important second messenger to control biofilm formation. In *P. aeruginosa*, there are 41 genes that are predicted to participate in the making and breaking this dinucleotide. A major missing information in this field is the systemic expression profile of those genes in response to changing environment. Toward this goal, we constructed a promoter-*gfp* transcriptional fusion reporters’ library that consists of 41 reporter plasmids, each of which contains a promoter of corresponding c-di-GMP metabolism-related genes in *P. aeruginosa*. This library provides a helpful tool to understand the complex regulation network related to c-di-GMP and to discover potential therapeutic targets.

## Introduction

Biofilms are termed as communities of microorganisms embedded in extracellular polymeric substances (EPS) (1). Biofilms can form on biotic and abiotic surfaces and exist as non-surface-attached aggregates(2-4), which protect microorganisms from harsh conditions and are a key feature of chronic and persistent infections(5). The second messenger *bis*-(3′-5′)-cyclic guanosine monophosphate (c-di-GMP) plays an important role in biofilm formation (6, 7). In general, a high level of c-di-GMP stimulates the biosynthesis of adhesins and exopolysaccharides to promote biofilm formation, whereas low level of c-di-GMP is associated with an increase in motility and virulence (8, 9). C-di-GMP is synthesized from two GTP molecules by diguanylate cyclases (DGC) that usually harbor a conserved GGDEF domain and is hydrolyzed by c-di-GMP specific phosphodiesterases (PDE) with conserved EAL or HD-GYP domains, which degrade c-di-GMP to pGpG(8). In some bacteria, such as *Pseudomonas aeruginosa*, there are multiple genes that were predicted to have a conserved GGDEF domain and/or EAL or HD-GYP domains(10). It remains largely unclear about the systemic expression pattern of those genes due to lack of available tools.

*P. aeruginosa* is a Gram-negative gamma-proteobacterium that can grow in diverse habitats and acts as an opportunistic pathogen over a wide range of hosts including human, animals and plants(11). The success of *P. aeruginosa* infections relies on both the production of acute virulence factors and the ability to form biofilms (12). This opportunistic pathogen has become a model bacterium for biofilm research. There are 41 genes in *P. aeruginosa*, whose encoding proteins were predicted to be involved in the synthesis and degradation of c-di-GMP, including 17 different proteins with a GGDEF domain, 8 with a EAL or HD-GYP domain, and 16 with both types of domains (10). Except for PA0290, PA0338, PA1851, PA2072, PA2572, PA3258, PA4396 and PA5442, the other 33 genes were functional characterized to encode a DGC or PDE(13-44). The enzymes with both GGDEF and EAL domains, are usually have only one catalytic activity, either PDE or DGC activity. However, enzymes encoded by PA0861 (*rbdA*), PA1727 (*mucR*) or PA4601 (*morA*) were found to conditionally switch between the two activities (35, 45-48). Intracellular c-di-GMP level in *P. aeruginosa* relies on the expression of those 41 <u>c</u>-di-GMP <u>m</u>etabolisms-related (CMR) genes. However, it remains a mystery about how *P. aeruginosa* controls the expression of those c-di-GMP metabolism enzymes.

*P. aeruginosa* is a ubiquitous microorganism, which is able to survive in a variety of environments. Its’ optimum growth temperature is 37°C. Growth occurs at temperatures as high as 42°C, but not at 4°C (49, 50). It has been reported that temperatures regulate biofilm formation of *P. aeruginosa* and the temperature-dependent changes in biofilm formation might be mediated by c-di-GMP (51). Henrik Almblad et al. have discovered a temperature-regulated DGC that coordinates temperature-dependent biofilm formation, motility, and virulence factor expression in *P. aeruginosa* CF39S, a strain isolated from the sputum of patients with chronic pulmonary cystic fibrosis (CF) (52, 53). Many environmental factors (such as oxygen levels, nitric oxide, iron, and nutrients etc.) and small chemical compounds, have been reported to trigger biofilm dispersion(35, 54-56). Studies have shown that sub-inhibitory concentrations of antibiotics can induce biofilm formation of *P. aeruginosa* (40, 57, 58). A systemic tool will be a great help for a deep understanding about how *P. aeruginosa* regulates the 41 CMR genes in response to various environmental temperatures, signals and antibiotics. A few reports have focused on the comprehensive and systemic features of those c-di-GMP related genes or proteins in bacteria (15, 17, 59-62), while those studies are mainly focused on the functions of genes or enzymes. Therefore, a systemic transcriptional profile of the 41 CMR genes in *P. aeruginosa* remains lacking.

In this study, we provide a promoter-*gfp* transcriptional fusion reporters’ library of the 41 genes that related to c-di-GMP metabolisms in *P. aeruginosa* for systemic transcriptional profile analysis. The plasmid pProbe-AT’ was selected as the vector to construct promoter-*gfp* transcriptional fusions because owing to its a broad-host range, high stability without antibiotic selection, and low background levels of expression in multiple taxa (63). More importantly, this promotor-probe vector is able to detect weak or moderate promoters. We have tested this CMR genes’ promoter-*gfp* transcriptional fusion reporters’ library in *P. aeruginosa* by different temperatures, growth media, and antibiotics. Each condition did exhibit a distinctive expression profile of CMR genes. Meanwhile, our results also revealed that some genes’ expression can be enhanced by a certain condition, which would provide the novel targets for the investigation and therapeutic on *P. aeruginosa* in the future.

## Material and methods

### Bacterial strains and growth conditions

*P. aeruginosa* strain PAO1 was cultured in Luria-Bertani (LB) medium (64), or Jensen’s medium(65), or M63 medium (66), or Artificial Sputum Medium (ASM) (67). *Escherichia coli* DH5α (Tsingke Biotechnology Co., Beijing, China) was inoculated in LB medium at 37°C, 200 rpm. The chemical composition of all media used in this study is shown in Table S1. When necessary, antibiotics were added at the following final concentrations: 300 μg/mL carbenicillin for *P. aeruginosa*; 100 μg /mL ampicillin for *E. coli*. The sub-inhibitory concentrations of antibiotics was applied as 30 μg/mL of erythromycin (68), or 2 μg/mL of azithromycin (69), or 0.3 μg/mL of tobramycin (40), or 0.0625 μg/mL of ciprofloxacin (70).

### Construction of promoter-*gfp* transcriptional fusion reporter plasmid

The plasmids constructed in this study are shown in Figure 1. The pProbe-AT’ (Ap^r^) was used as a vector for promoter-Gfp transcriptional fusions (63). The websites (http://www.phisite.org/ and https://services.healthtech.dtu.dk/ and https://www.fruitfly.org/seq_tools/promoter.html) were used to predict the promoter region and RBS (ribosome binding site) region of target genes (71-73). The information of cloned promoter regions and their corresponding genes and PA# was also listed in Figure 1. The cloned promoter sequences consist of the entire intergenic region, together with 0-500 bp upstream genes and 0-138 bp of the coding region of corresponding ORFs (<u>o</u>pen reading frames). All promoters were amplified from the genome DNA of *P. aeruginosa* PAO1 and then cloned into the *Eco*RI*-Bam*HI sites of pProbe-AT’. The cloned regions in each corresponding plasmid were all confirmed by sequencing. The promoter-Gfp fusion plasmid was introduced into the *P. aeruginosa* strain by chemical transformation in order to determine the promoter’s activity in *P. aeruginosa*.

**Fig 1.**
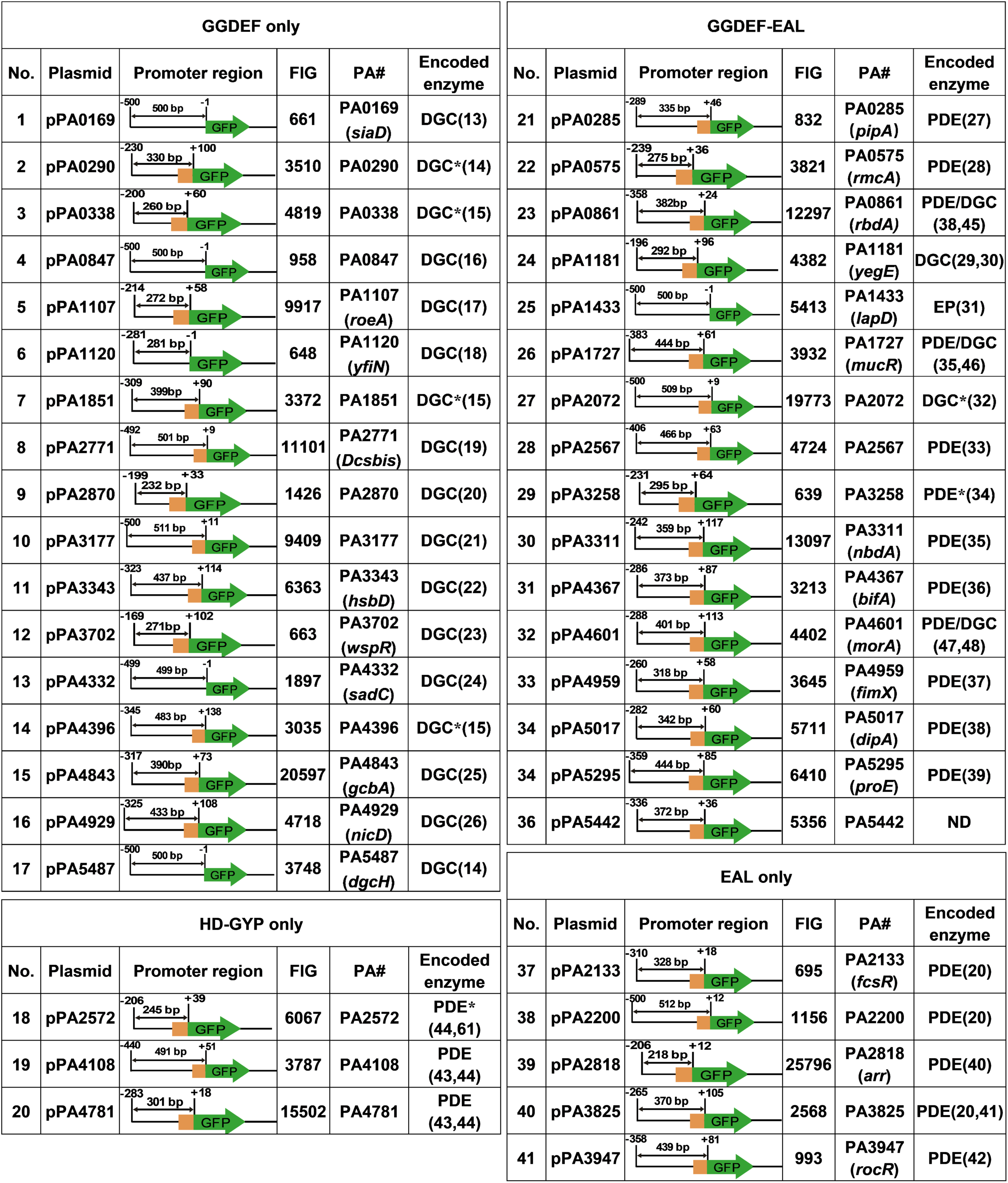
The information of 41 plasmids in the library of <u>c</u>-di-GMP <u>m</u>etabolism-related (CMR) <u>g</u>enes’ promoter-Gfp transcriptional fusion reporters and corresponding genes. +1, the first base of the start codon ; ND, not determined; F.I., fluorescence intensity; PA#, corresponding locus number in genome of *P. aeruginosa* PAO1 ; *, the function is not fully identified; FIG, <u>F</u>luorescence intensity of <u>G</u>FP. Shown is the average fluorescence intensity value per OD_600_ of the corresponding sample grown in Jensen’s medium at 37°C for 24 hours.

### Isolation of total RNA and qRT-PCR

PAO1 was cultivated in Jensen’s medium at 37°C, 200 rpm until the OD_600_ reached 3.0, at which point 1 mL of cells were harvested and RNA samples were extracted using the RNAprep Pure Bacteria Kit (Tiangen Co., Beijing, China). Total cDNA was generated from 500 ng of total template RNA using the HiScript III All-in-one RT SuperMix Perfect for qPCR (Vazyme Co., Nanjing, China). The ChamQ Universal SYBR qPCR Master Mix (Vazyme Co., Nanjing, China) and gene-specific primers were used to perform quantitative reverse transcription-PCR (qRT-PCR) on 5 ng/μL cDNA in a Real-Time PCR System (Applied Biosystems ViiA 7, USA). Table S2 lists the primers used in qRT-PCR. Real-time PCR data were analyzed, validated, and calculated according to the instructions of the manufacturer. All samples were normalized to constitutively produced *rpsL* transcripts.

### Measurement of fluorescence intensity

To determine the fluorescence intensity of the promoter, The overnight cultures of the plasmid-carrying strains grown in Jensen’s medium with carbenicillin or ampicillin, were inoculated at 1% into 200 μL of Jensen’s medium (unless otherwise specified) containing carbenicillin or ampicillin in 96-well plates (NEST Co., Wuxi, China) and grow with shaking at 700 rpm in a constant temperature microplate shaker (MIULAB Co., Hangzhou, China) at 37°C (unless other specific temperature was required) until the time for measurement (8 hours for exponential phase culture for 24 hours for stationary phase culture). Fluorescence of GFP was measured on a Synergy H4 hybrid reader (BioTek, USA) at excitation wavelength at 488 nm and emission wavelength of 520 nm with gain 50. Biomass was determined at 600 nm with gain 20 as scattered light (14). Fluorescence signal value was normalized to OD_600_, and fluorescence derived from pPROBE-AT’ was subtracted to account for background fluorescence. All experiments were performed with a minimum of three biological triplicates. Statistically significant differences were determined using a two-tailed Student’s t-test (*P* < 0.05).

## Results

### Construction of a promoter-Gfp transcriptional fusion reporters’ library for monitoring transcriptional profile of 41 CMR genes in *P. aeruginosa*

The promoter regions of 41 genes that are (or are predicted to be) involved in c-di-GMP metabolism in *P. aeruginosa* PAO1 were cloned into the vector pPROBE-AT’ respectively, resulting in a library of promoter-Gfp transcriptional fusion reporter plasmids (Figure 1), which can monitor the expression of the corresponding genes in *P. aeruginosa*.

To evaluate whether the cloned promoter regions were functional, each promoter-Gfp fusion plasmid was transformed into PAO1, respectively. Their fluorescence intensity was measured in Jensen’s medium at 37°C. The fluorescence intensity of these reporters was varied from hundreds to over 10 thousand, indicating that all promoters can initiate the transcription of *gfp*. Among the 41 genes, the promoter of PA2818 (*arr*) showed the highest fluorescence intensity (Figure 2A). PA2818 encodes a PDE named Arr, which is responsible for aminoglycoside antibiotics-induced biofilm formation (40). Moreover, fluorescence intensity of the other five genes, namely PA4843 (*gcbA)*, PA4781, PA0861 (*rbdA*), PA2072, and PA3311 (*nbdA*), is also above 10,000. Among these five genes, PA4843 encodes a DGC GcbA (25), PA4781 and PA3311 (*nbdA*) both encode functional PDE, PA0861 (*rbdA*) has intrinsic GTP-stimulated PDE activity as well as DGC activity (38, 45), while the function of PA2072 encoding enzyme has not determined so far (25, 74). Except for PA2818 (*arr*), the fluorescence intensity of stationary phase culture is higher than that of the exponential phase culture (Figure 2A), which is consistent with the stability of Gfp. According to previous publications (15, 75), the two key DGCs, SadC and SiaD showed stronger effects on intracellular c-di-GMP concentration than other DGCs of *P. aeruginosa*, yet our results indicated their transcriptional level was relatively low among DGCs (Figure 1). We then selected five genes, PA0169 (*siaD*), PA2818 (*arr*), PA4332 (*sadC*), PA4601 (*morA*), and PA4843 (*gcbA)*, to detect their transcription by qRT-PCR. As shown in Figure S1, the transcriptional level of these genes is consistent with the results of promoter-Gfp, indicating that the library can be used to monitor the expression pattern of the corresponding genes. Given that pPROBE-AT’ is a stable plasmid, we compared fluorescence intensity of the 41 plasmids with and without the addition of antibiotics. Without antibiotics, the fluorescence intensity of reporters showed some decrease but not significant, except for that of a few genes (especially those with fluorescence intensity over 10-thousand), suggesting that the library could be used without antibiotics in growth medium (Figure 2A) or *in vivo* study.

**Fig 2.**
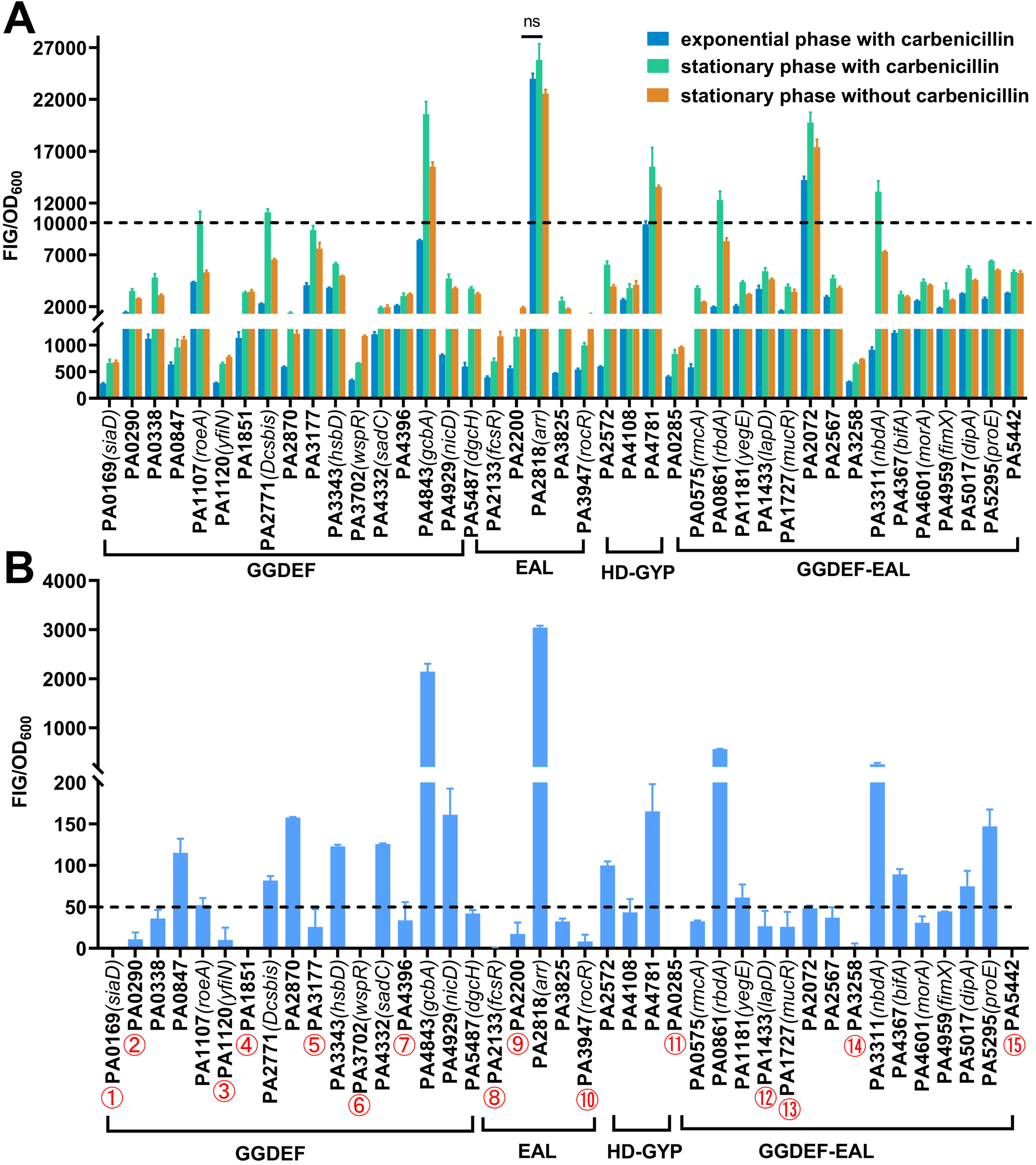
The fluorescence intensity of CMR’ gene promoter-Gfp fusion reporter library in *P. aeruginosa* or *E. coli* and detecting the transcription of selected genes by qRT-PCR. FIG, <u>F</u>luorescence intensity of <u>G</u>FP. A, the fluorescence intensity CMR’ promoter-Gfp fusion reporter in *P. aeruginosa* cultures at exponential phase with carbenicillin, stationary phase with carbenicillin, and stationary phase without carbenicillin were examined and the fluorescence intensities was normalized by corresponding OD_600._ B, the fluorescence intensity of CMR’ promoter-Gfp fusion reporters in *E. coli*. the fluorescence intensity of CMR’ promoter-Gfp fusion reporters of *E. coli* stationary phase cultures. FIG, <u>F</u>luorescence intensity of <u>G</u>FP. The fluorescence intensities were normalized by corresponding OD_600._ Shown were the fluorescence intensities of stationary phase cultures normalized by corresponding OD_600_.

To test whether the 41 promoters could function in *E. coli*, the fluorescence intensity of promoter-Gfp fusion plasmids in *E. coli* strains were also measured under the same culture conditions. As shown in Figure 2B, the promoters of four genes, namely PA4843 (*gcbA)*, PA2818 (*arr*), PA0861 (*rbdA*), and PA3311 (*nbdA*), keep high activity in *E. coli*, while their fluorescence intensity are approximately 10-fold less than that in *P. aeruginosa*. The promoters of 15 genes (Figure 2B, marked with number) showed baseline level of fluorescence intensity, suggesting that these genes’ transcription might be *Pseudomonas*-specific. The other 22 genes’ promoters showed lower expression level (approximately ≥50) (Figure 2B).

### Effects of temperature on the transcriptional profile of CMR genes in *P. aeruginosa*

*P. aeruginosa* can live in diverse ecological niches. Therefore, we investigated whether temperature affected the transcription of genes that were involved in c-di-GMP metabolism in PAO1. The 41 strains containing corresponding plasmids were cultured at 25°C or 30°C, and their fluorescence intensity was compared to that of their corresponding culture grown at 37°C. As shown in Figure 3, PA2567 exhibited a higher expression at both 25°C and 30°C, which is the only gene that has high-level of transcription at 25°C, suggesting that this gene might be important for biofilm formation at lower temperatures. Overall, the transcription levels of most genes were relatively low at 25°C or 30°C compared to that of 37°C, while there are 10 genes, namely PA0338, PA3177, PA3343 (*hsbD*), PA4843 (*gcbA*), PA2818 (*arr*), PA1433 (*lapD*), PA4367 (*bifA*), PA4601 (*morA*), PA4959 (*fimX*), and PA5295 (*proE*), have similar transcriptional levels at three tested temperatures (Figure 3, marked with red number). Among these 10 genes, PA3177 and PA3343 (*hsbD*) encode DGCs (21, 22), PA2818 (*arr*), PA4367 (*bifA*), PA4959 (*fimX*) and PA5295 (*proE*) encode PDEs (36, 37, 39, 40), while PA4601 (*morA*) functions as both DGC and PDE. Furthermore, MorA is conserved among diverse *Pseudomonas* species (47, 48). In addition, 7 genes exhibited lower-level expression at 25°C, yet had similar expression level at 30°C when compared to the corresponding samples at 37 °C (Figure 3, marked with black number). Taken together, our results showed that lower temperature generally reduced the expression of genes involved in c-di-GMP metabolism with some exceptions, and suggesting that the gene with high level transcription at lower temperature might be important for the survival of *P. aeruginosa* in environment.

**Fig 3.**
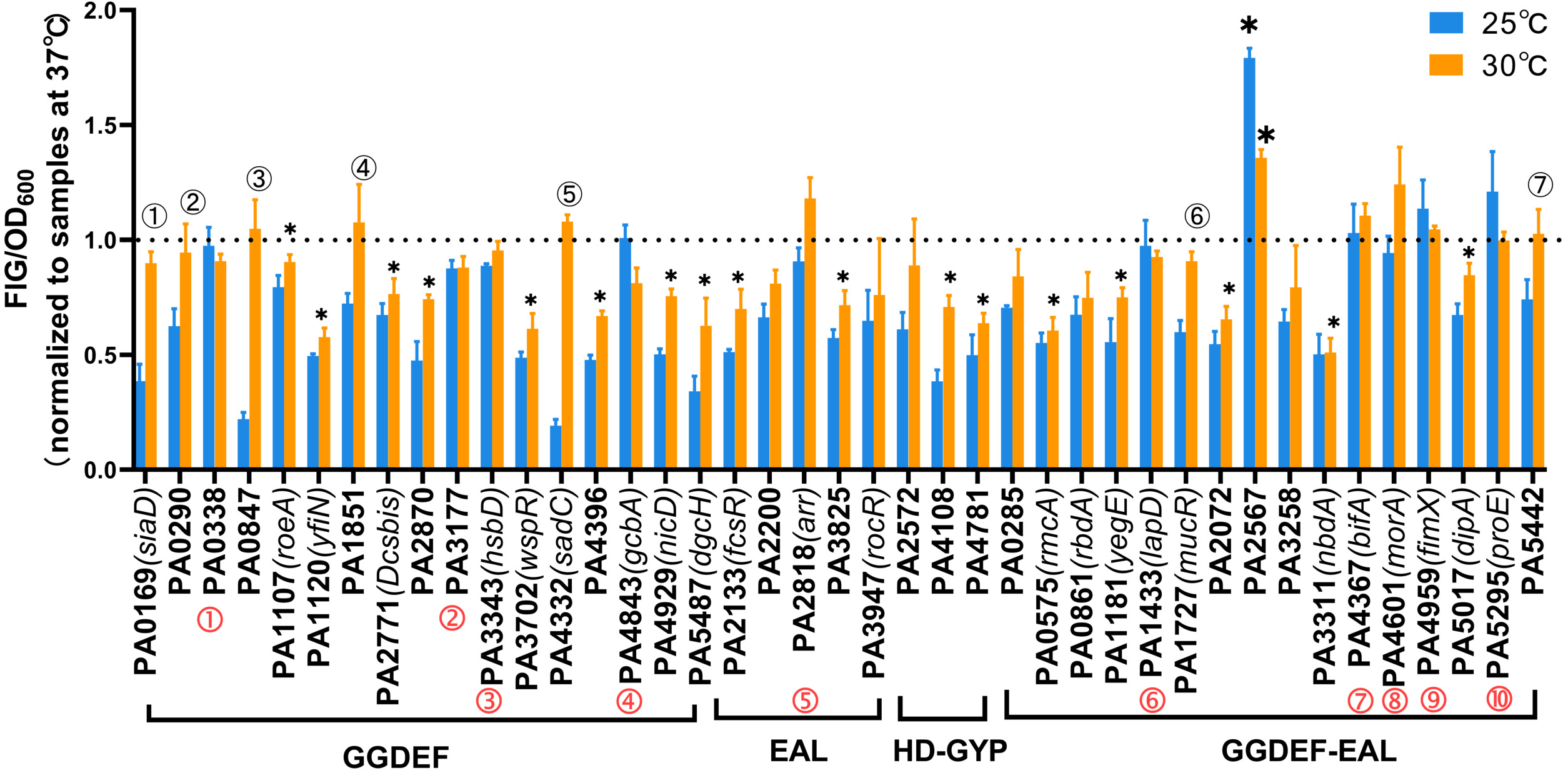
Effects of different temperatures on the promoter activity of each CMR gene in *P. aeruginosa* PAO1. Blue column, 25°C; orange column, 30°C. The value was normalized to the fluorescence intensity per OD_600_ of the corresponding strain grown in Jensen’s medium at 37°C; Red numbers indicated the genes that their promoter activity did not change at these three temperatures; Black numbers marked the genes that their promoter activity at 30°C were similar as that at 37°C; Small asterisk, significantly decreased; Large asterisk, significantly increased. *, *P* < 0.05. Error bars were calculated from three independent experiments.

### Effects of different media on the transcriptional profile of CMR genes in *P. aeruginosa*

To investigate whether nutrient affects the transcriptome of genes related to intracellular c-di-GMP metabolism, we examined 41 genes’ expression in four media, minimal medium M63, chemical defined Jensen’s medium, rich medium LB, and artificial sputum media (ASM), respectively (see Table S1 for chemical composition). The fluorescence intensity of each promoter-Gfp fusion reporter was normalized to its corresponding value in Jensen’s medium grown at 37°C (Figure 4).

**Fig 4.**
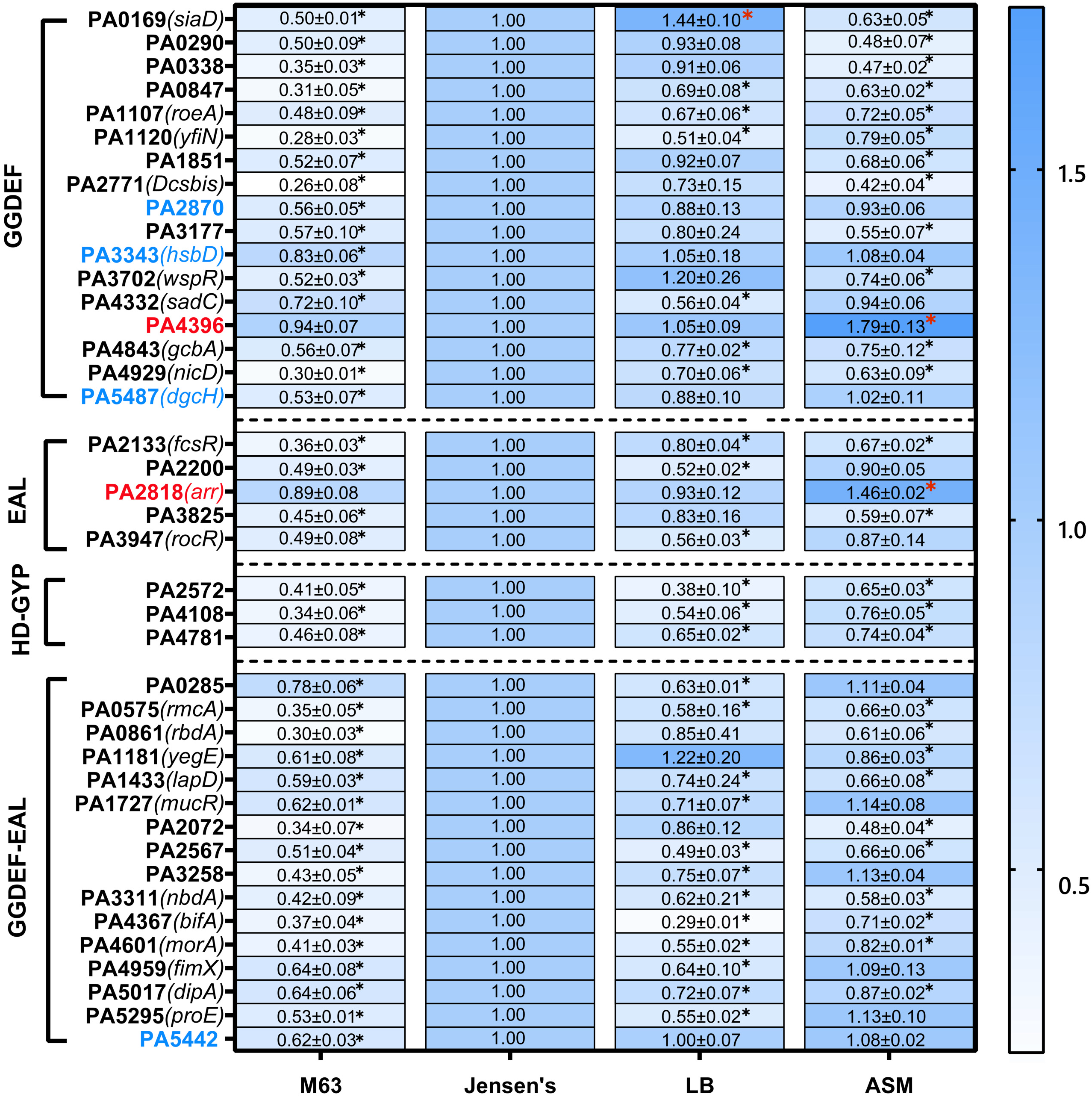
Effects of different growth media on the promoter activity or expression of CMR’ genes in *P. aeruginosa* PAO1 background. The value was normalized to the fluorescence intensity per OD_600_ of the corresponding strain grown in Jensen’s medium at 37°C. Black asterisks represent significantly reduced; Red asterisks represent significantly elevated. Genes highlighted in red were significantly elevated in ASM with no change in the other media; Genes highlighted in blue, significantly reduced in M63 with no change in the other media. Shown were averages with standard division calculated from three independent experiments. *, *P* < 0.05.

The promoter activities (or expression levels of corresponding gene) significantly decreased in M63 minimal medium compared to Jensen’s medium, except for PA2818 (*arr*) and PA4396, which kept the same level in M63 as in Jensen’s medium (Figure 4, marked in red). In the LB medium, most genes exhibited similar (16 out of 41 genes) expression level or reduced level (24 out of 41 genes) compared to that in Jensen’s medium, while PA0169 (*siaD*), as an important DGC, showed a significantly increased expression (Figure 4, indicated with a red star). However, in ASM, PA2818 (*arr*) and PA4396 showed increased expression (Figure 4, marked by a red star), 12 out of 41 genes (PA2870, PA3343, PA4332, PA5487, PA2200, PA3947, PA0285, PA1727, PA3258, PA4959, PA5295 and PA5442) have similar expression level, and the other 27 genes’ expression decreased compared to that in Jensen’s medium.

Strikingly, the promoters of PA2818 (*arr*) and PA4396 exhibited the highest activity in the Jensen’s medium (Figure 1), which remained at the same level in either M63 or LB, and even higher level in ASM. These results suggested the importance of these two genes. In addition, there are four genes, PA2870, PA3343 (*hsbD*), PA5487 (*dgcH*) and PA5442 (marked with blue color), whose promoter activities were only reduced in M63 and kept the same level in the other three media. PA5487 (*dgcH*) was reported to be a conserved DGC that its expression is highly invariable, which is consistent with our results (14).

### Effects of sub-inhibitory concentrations of antibiotics on the expression pattern of CMR genes

Previous reports showed that sub-inhibitory concentrations of aminoglycoside antibiotics induced the biofilm formation of *P. aeruginosa* (40). We then examined the effect on our reporter’ library by three types of antibiotics at sub-inhibitory concentrations, including aminoglycoside antibiotics tobramycin, fluoroquinolone antibiotics ciprofloxacin, macrolide antibiotics erythromycin and azithromycin, respectively. Our results showed that tobramycin increased the expression of PA2818 (*arr*) (Figure 5A), which is consistent with the previous report about the contribution of PA2818 on the tobramycin-induced biofilm formation (40). The expression of two genes, PA5295 (*proE*) and PA5442 are also induced under sub-inhibitory concentrations of tobramycin (Figure 5A). ProE is a very active PDE with high enzymatic activity in the degradation of c-di-GMP and plays an important role in regulating EPS production in *P. aeruginosa* PAO1 (39), and the function of PA5442 is not determined. The transcription of 7 genes (PA1107, PA4332, PA4396, PA4781, PA4601, PA4959, PA5017) were not changed by tobramycin. The other 28 genes’ expression were significantly reduced in the presence of tobramycin.

**Fig 5.**
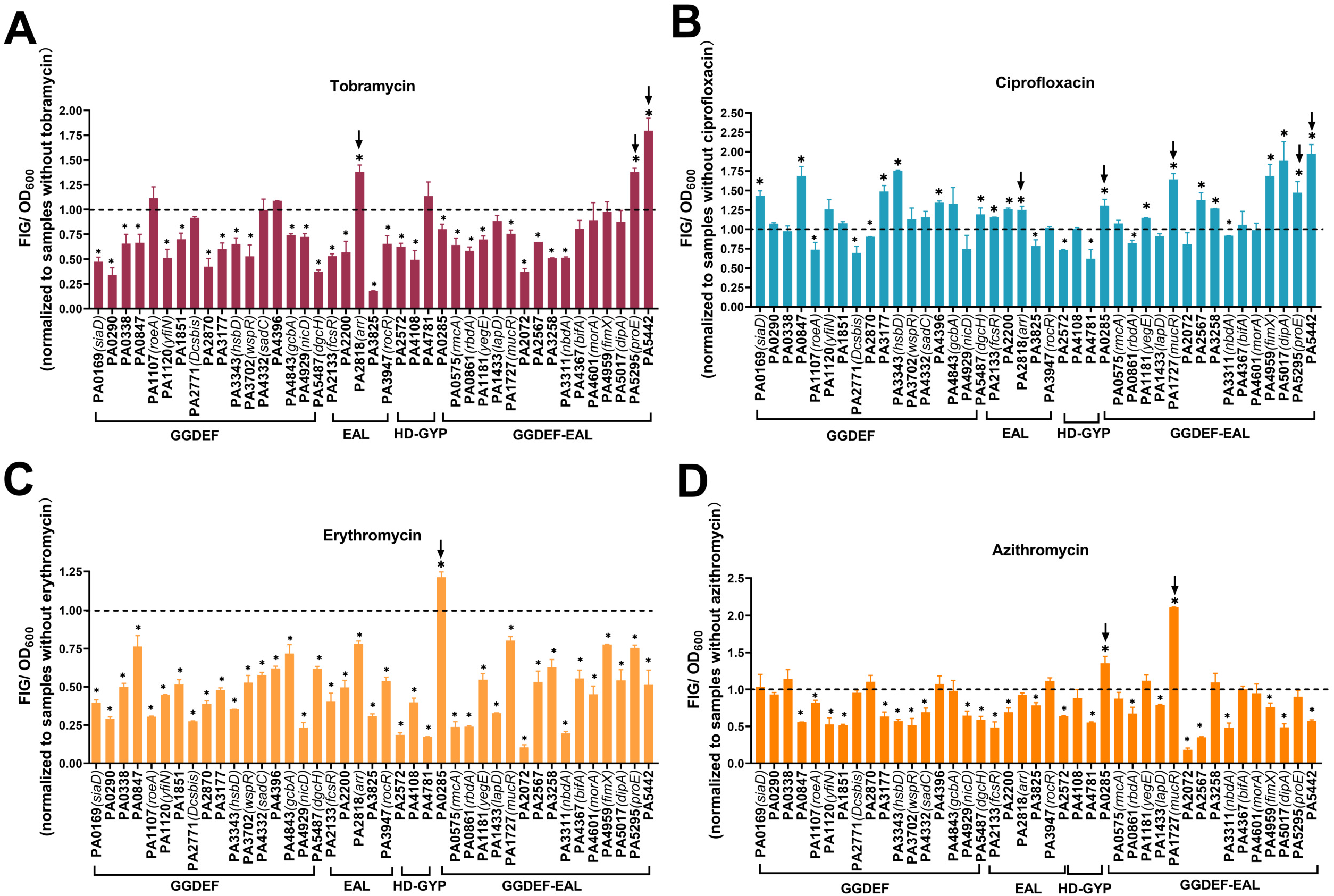
Effects of sub-inhibitory concentrations of antibiotics on the promoter activity or expression of CMR’ genes in *P. aeruginosa* PAO1. A-D, the results of tobramycin, ciprofloxacin, erythromycin, and azithromycin treatment, respectively. Each value was normalized to the corresponding sample without antibiotic treatment. The genes with promoter activity significantly elevated in more than 2 antibiotics were indicated with arrows; Small asterisks marked the genes with significantly reduced level and large asterisk indicated those with significantly increased level. Error bars were calculated from three independent experiments. *, *P* < 0.05.

Among the four antibiotics, ciprofloxacin induces the expression of a large number of genes, as shown in Figure 5B, 18 genes’ expression was significantly elevated, while 8 genes’ expression was decreased, and 15 genes’ expression showed no changes. It is worth noting that expression of PA2818 (*arr*), PA5295 (*proE*), and PA5442, were enhanced by both tobramycin and ciprofloxacin.

For erythromycin and azithromycin, they both repressed transcription of most c-di-GMP metabolism genes, only a few genes’ expression is enhanced (Figure 5C). PA0285 (*pipA*) is the only gene whose transcription is enhanced by both erythromycin and azithromycin. The transcription of PA1727 (*mucR*) is enhanced by azithromycin, but reduced by erythromycin.

## Discussion

C-di-GMP is an important second messenger involved in bacterial switching from motile to sessile lifestyles. It is critical for bacteria to control the intracellular c-di-GMP level in response to changing environments. The opportunistic pathogen *P. aeruginosa* can live in diverse ecological niches. Consistently, it has 41 genes that are predicted to encode proteins for the synthesis or degradation of c-di-GMP. Extensive studies have been done to study their functions in respect to biofilm formation and motilities. However, it remains a mystery about when and where those genes will be transcribed and there is still a paucity of information concerning the systemic expression pattern of those genes. In this study, we have constructed a promoter-*gfp* transcriptional fusion reporters’ library for systemic investigating the transcription or regulation of the 41 c-di-GMP’s metabolism-related genes.

We have examined this <u>c</u>-di-GMP <u>m</u>etabolism-related (CMR) genes’promoter-Gfp fusion library by different growth conditions, including different media, temperatures, or antibiotics. Each condition does affect different genes’ transcription and some genes’ transcription are induced in a typical condition. For example, the expression of PA2567 is enhanced by lower temperature (25°C>30°C>37°C). The sub-inhibitory concentrations of tobramycin induce the transcription of the three genes: PA2818 (*arr*), PA5295 (*proE*) and PA5442 respectively. A previous study has showed that PA2818 (*arr*) is important for biofilm formation induced by aminoglycoside antibiotics(40), whereas the functions of PA5295(*proE*) and PA5442 have not been linked with antibiotics. Our results have shown that this library is not only helpful for studying the systemic expression pattern of CMR genes, but also can reveal the new roles of those genes.

It is worth noting that PA2818 (*arr*) has the highest transcription among the 41 CMR genes. Moreover, its transcript is consistently stable in several tested conditions and elevated when grown in artificial sputum media or in the presence of sub-inhibitory concentrations of aminoglycoside antibiotics. These results suggested that PA2818 (*arr*) might play a key role in biofilm-related persistent infections caused by *P. aeruginosa*. Given that *P. aeruginosa* can cause life-threatening lung infections in cystic fibrosis patients, our results have also provided a therapeutic target.

The promoter-probe vector pProbe-AT’ used in this study is a broad-host range vector, thus we have examined the promotor’s activity in both *P. aeruginosa* and *E. coli*. The promotors of 26 genes can initiate the transcription of *gfp* in *E. coli*., suggesting these promoters might function in Gram-negative bacteria. Therefore, this promoter-*gfp* library also provide a library of bio-bricks for synthetic biology.

In summary, the CMR gene’ promoter-*gfp* transcriptional fusion reporters’ library we have constructed in this study is a versatile and helpful tool. The library can be used in the following aspects: i, investigating the systemic expression pattern and regulatory network of CMR genes in *P. aeruginosa*; ii, determining the regulatory mechanism of any factor that affects intracellular c-di-GMP in *P. aeruginosa*; iii, discovering compounds or drugs that target at the intracellular c-di-GMP of *P. aeruginosa*; iv, elucidating the role of CMR genes and their regulated pathways; V, a promoter library for future applications in synthetic biology.

## Acknowledgements

Authors thank Dr. Xi Liu in the Institute of Microbiology, Chinese Academy of Sciences, for critical reading on manuscript, Zhenyu Zhang and Qi Zhou for help in figure making. This work is supported by the National Key R&D Program of China (2021YFA0909500, 2019YFC804104, and 2019YFA0905501), the National Natural Science Foundation of China (91951204, 32070033). The funders had no role in the study design, data collection and interpretation, or the decision to submit the work for publication.

## Figure legends

Figure S1. The relative transcriptional level of PA0169 (*siaD*), PA4332 *(sadC*), PA4601 (*morA*), PA4843 (*gcbA*), and PA2818 (*arr*) to PAO1 was quantified by qRT-PCR. The house keeping gene *rpsL* was used as an endogenous control to normalize the quantification of the mRNA target. **, *P* < 0.01; ***, *P* < 0.001.

**Table S1.**
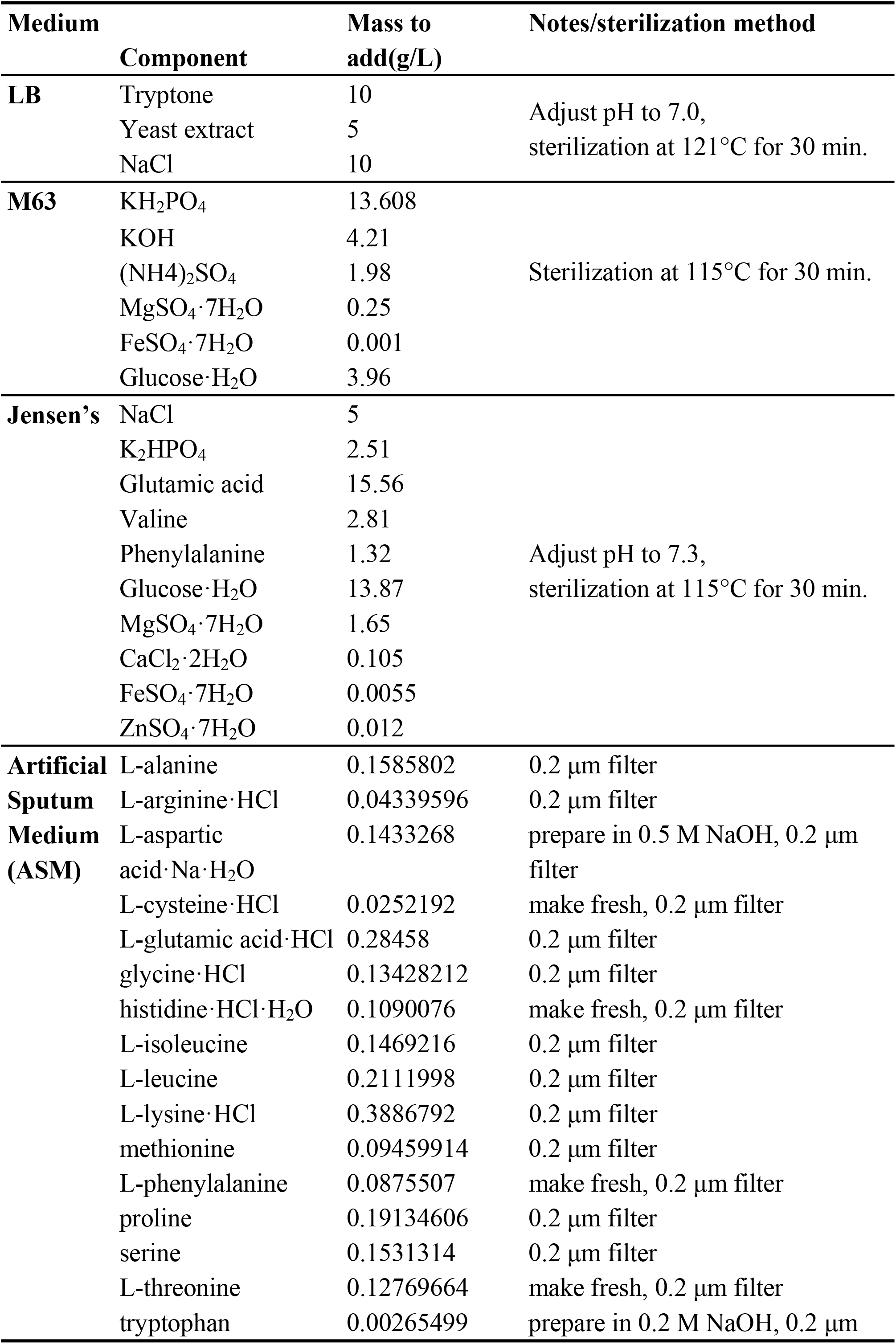
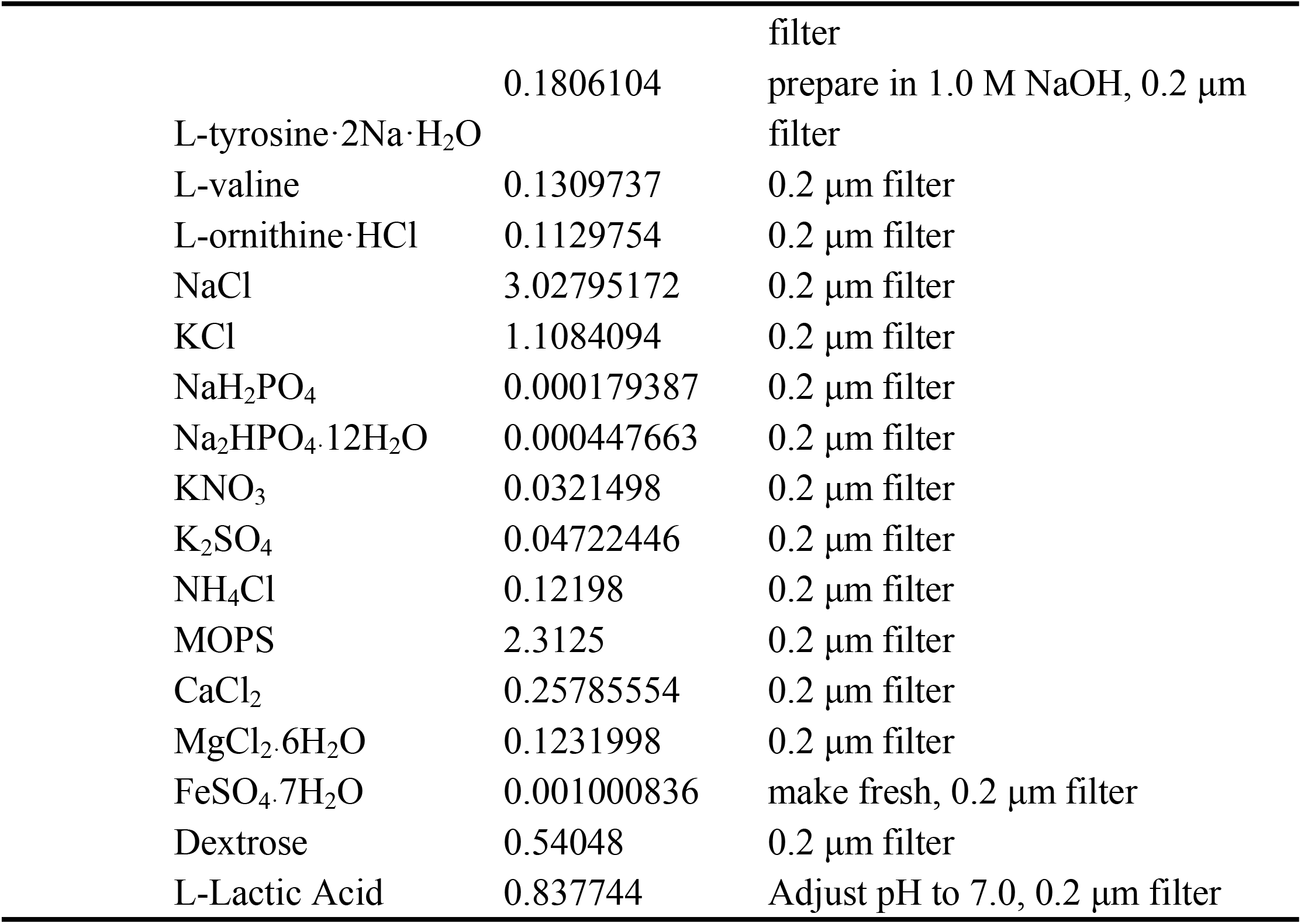
The chemical composition of media used in this study.

**Table S2.**
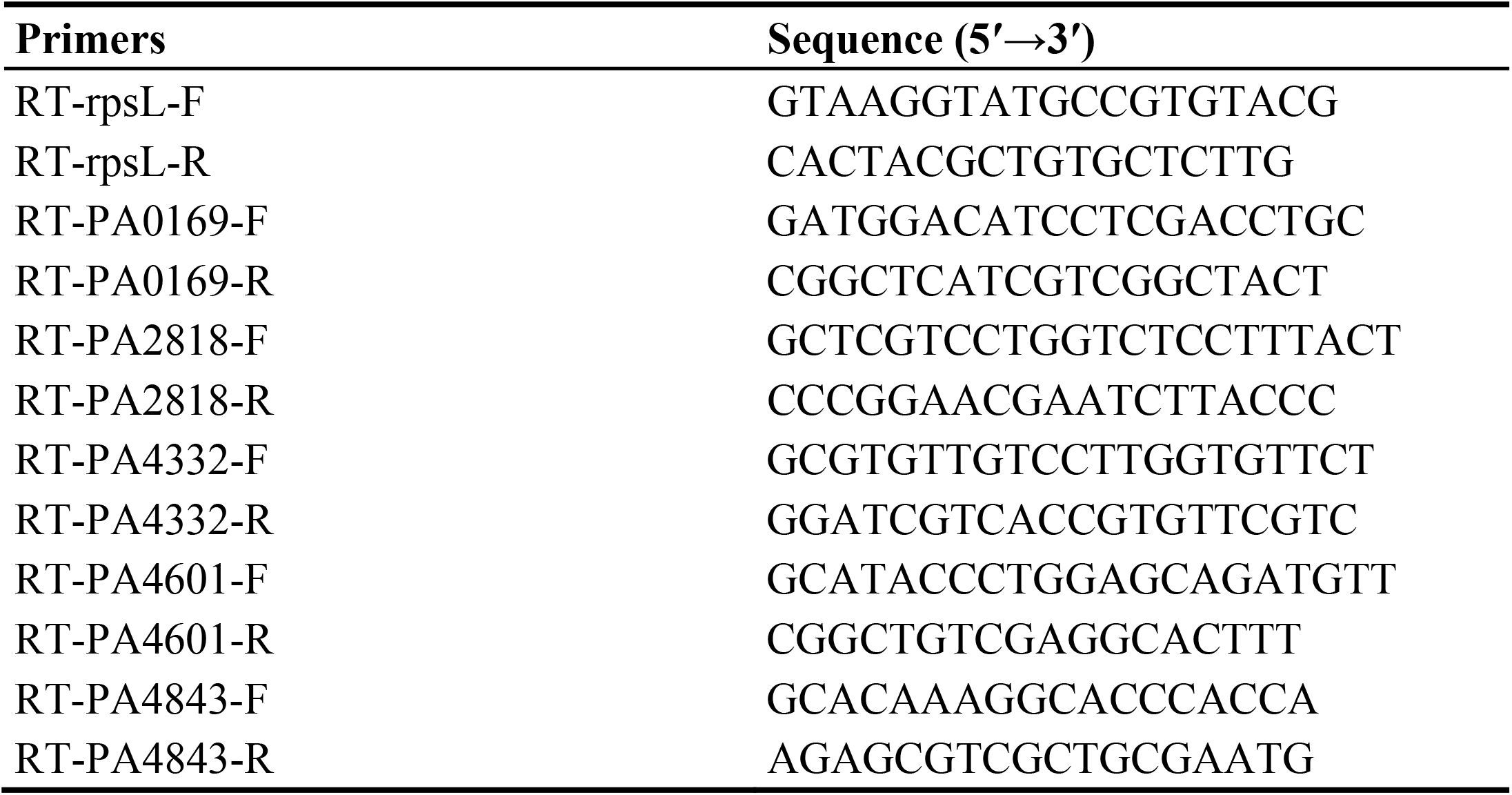
Primers used for qRT-PCR in this study.

